# Nonequilibrium switching of segmental states can influence compaction of chromatin

**DOI:** 10.1101/2023.06.21.546028

**Authors:** Soudamini Sahoo, Sangram Kadam, Ranjith Padinhateeri, P. B. Sunil Kumar

**Author notes:** National Institute of Technology Rourkela, Odisha, 769008, India.

## Abstract

Knowledge about the dynamic nature of chromatin organization is essential to understand the regulation of processes like DNA transcription and repair. While most models assume protein organization and chemical states along chromatin as static, experiments have shown that these are dynamic and lead to the switching of chromatin segments between different physical states. To understand the implications of this inherent nonequilibrium switching, we present a diblock copolymer model of chromatin, with switching of its segmental states between two states, mimicking active/repressed or protein unbound/bound states. We show that competition between switching timescale *T*_*t*_, polymer relaxation timescale *τ*_*p*_, and segmental relaxation timescale *τ*_*s*_ can lead to non-trivial changes in chromatin organization, leading to changes in local compaction and contact probabilities. As a function of the switching timescale, the radius of gyration of chromatin shows a non-monotonic behavior with a prominent minimum when *T*_*t*_ ≈ *τ*_*p*_ and a maximum when *T*_*t*_ ≈ *τ*_*s*_. We find that polymers with a small segment length exhibit a more compact structure than those with larger segment lengths. We also find that the switching can lead to higher contact probability and better mixing of far-away segments. Our study also shows that the nature of the distribution of chromatin clusters varies widely as we change the switching rate.

**Significance statement:** Different cells in multicellular organisms have the same DNA but different functions. The function of any given cell type can be time-dependent. The current understanding is that differences in gene expression arising from local compaction and the probability for far-away regulatory segments to come in contact play an important role in establishing these differences. The necessary structural variations are achieved through a combination of changes in the chemical and physical states of chromatin regions. In this paper, we present a model for chromatin accounting for the dynamic switching of chromatin regions between different chemical and physical states. We demonstrate the implications of such switching in determining the local 3D structure of chromatin.

## I. INTRODUCTION

Chromatin is a long polymer made of DNA and proteins [1, 2]. Beyond the genetic code, the chromatin polymer encodes epigenetic information via the protein organization along the polymer, post-translational modifications (PTMs) on proteins, and the 3D organization of its contour [3–6]. Based on a number of experimental and theoretical studies, it is understood that chromatin structure is highly dynamic [7–10]. Apart from the thermal fluctuation of the polymer, the dynamicity also arises from the kinetics of epigenetic states, such as time-dependent changes in histone modifications and other protein organization [7–11]. The binding and dissociation of proteins also lead to local alterations in the chemical and physical states of the polymer [12–15]. In most of the cells, particularly stem cells, chromatin segments are known to switch between different epigenetic states exhibiting a bimodal distribution in the net amount of repressive/active epigenetic marks [16, 17]. These fluctuations have an impact on the way cells process information and make “decisions” [18–21]. For example, it is known that gene regions switch between transcribing and transiently inactive states, leading to bursts in mRNA production [22, 23].

The changes in epigenetic states and switching dynamics are known to be strongly coupled to the 3D organization of chromatin [22–24]. Recent experimental advances such as Hi-C and super-resolution imaging methods have made it possible to measure the 3D organization of chromatin [25]. The findings from these studies suggest that chromosomes are broadly organized into looped structures known as Topologically Associated Domains (TADs) [26, 27]. Subsequent experiments probing the contact map at higher resolution discovered the correlation between locations of chromatin-binding proteins (like CTCF and cohesin) and TAD boundaries, elucidating the role of the loop extrusion mechanism [28–31]. These investigations gave measurements of contact probability that two parts of the chromosomes along the contour are in contact as a function of the gnomic separation. Subsequent experiments advanced the field by not only obtaining the contact map but also by measuring the 3D distance between certain selected regions of chromatin using microscopy and techniques like FISH [14, 32–34]. The 3D organization of chromatin mediated by some set of proteins (for example, CTCF, LEFs, HP1) are widely studied. However, it is likely that there are other ingredients that can also simultaneously influence chromatin configurations in space and time. In this paper, we would discuss one such aspect wherein the chemical and physical states of local chromatin segments are changed in a non-equilibrium way.

The availability of experimental data on contact maps, 3D distances, and TAD organization of chromatin led to two sets of theoretical studies using concepts from polymer physics. One set examined the nature of intra-chromosomal interactions that can generate experimentally observed contact probabilities and 3D distances [26, 35–51]. These include both gene-specific models and generic models that might be applicable to a variety of gene locations. The considered models include homogeneous polymers with various kinds of interchromatin interactions and block copolymers that are heterogeneous [36, 42, 47, 52–55].

While almost all of the above theoretical studies investigated the equilibrium organization or dynamics that lead to the equilibrium of passive polymers, there have been a set of nonequilibrium studies examining the role of loop extrusion, or the interplay between loop extrusion and intra-chromatin interactions resulting in the experimentally observed 3D organization [29, 53, 56– 65]. The effect of enzymes such as topoisomerase in the organization of chromatin was studied recently [66]. These consist of the second set of studies. Ramakrishnan *et al*. looked at a semiflexible polymer with active stress fluctuations and found that activity drives the chromatin to nonequilibrium steady-state conformations exhibiting statistics consistent with experimental observations of intra-chromosomal contact probabilities and chromosomal compaction [59]. Recently there have been a few models examining the kinetics of histone modifications [67, 68]. Katava et al. studied the spreading of histone modifications around a nucleation site coupled with polymer dynamics[67]. Sood et al. proposed a model that includes bi-stable histone modifications to study the interplay between different time scales arising from chemical state switching and confirmation changes [68].

There are multiple ways through which the local regions of chromatin polymer can switch between different chemical and physical states. For example, the assembly and disassembly of nucleosomes will make the polymer compact/extended. Similarly, repression/activation of genes will make the gene regions compact/extended. In all the cases, changes in the inter and intra-segmental interactions affect the steady-state configuration of the chromatin. In the past, the effect of intra-segmental and inter-segmental interactions on the chromatin structure are studied only with a static distribution of segmental states along the chromatin polymer [40, 53, 63].

Here we study a model that allows us to investigate the interplay between the switching in chromatin states and the 3D organization of chromatin. We consider a diblock copolymer with blocks that can actively switch between solvo-philic and solvo-phobic states leading to their corresponding physical states. We systematically vary the length of the segments and explore the interplay between switching time and different relaxation times of the polymer. We show that, at steady state, a) the nonequilibrium switching can lead to a non-monotonic variation in the radius of gyration of the polymer as a function of the switching time, b) the contact probability at small separations increases monotonically with switching time but has a non-monotonic dependence for large separations, and c) while polymer could have similar overall compaction for different switching rates, the local organization of the polymer is very different as could be inferred from the cluster distribution.

## II. MODEL AND METHODS

We model chromatin as a flexible linear diblock copolymer made of *N*_*p*_ beads connected with springs, having segments of length *N*_*s*_ randomly arranged along the polymer (see Fig. 1(a)). A segment can have multiple physical interpretations. For example, each segment can be considered as a nucleosome, and the two states of the segments in the diblock copolymer would correspond to the bound and unbound states of the nucleosomes (see the details below). This would mean that, for *N*_*s*_ = 4, each bead corresponds to approximately 50 bp. Alternatively, for higher segment length *N*_*s*_ = 16, we could take each segment to represent a gene or promoter region with a size of the order of 5 kb. In this case, for *N*_*s*_ = 16, one bead will be equivalent to one nucleosome. To mimic the binding and dissociation of nucleosomes and the associated interactions between nucleosomes, the segments are assigned to have either a “solvo-philic” (nucleosome unbound) or “solvo-phobic” (nucleosome bound) nature, represented by the variable *ϕ* = 1 and *ϕ* = −1, respectively. Solvo-philic beads behave like a polymer in a good solvent [69, 70]. On the contrary, solvo-phobic beads behave similarly to a polymer in a bad solvent [69, 70]. The segments can switch between these two states in a nonequilibrium way. We use Monte Carlo simulations to sample the conformations of the polymer chain.

**FIG. 1.**
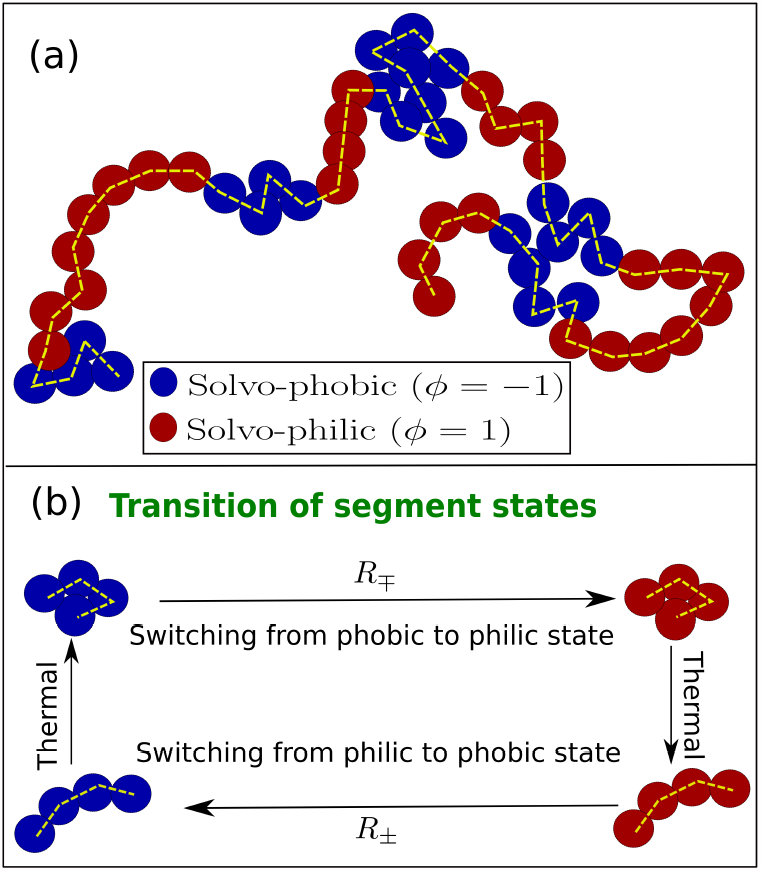
(a) Schematic representation of a block copolymer with solvo-phobic (*ϕ* = −1) and solvo-philic (*ϕ* = 1) segments with *N*_*s*_ = 4. The polymer backbone is shown as the yellow dashed line. (b) Schematic of the loop of segmental state transition from *ϕ* = −1 to *ϕ* = −1 and vice versa. See supplementary material, Fig. S1 for an analysis of the Kolmogorov condition and the corresponding equilibrium transition rules.

The neighboring beads of the copolymer are connected through finite extensible nonlinear elastic (FENE) springs having the energy of the form,

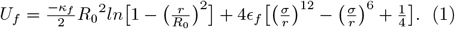

Here, *κ*_*f*_ is the spring constant, *r* = |*r*_*i*_ − *r*_*i*+1_| is the separation between the consecutive beads, and *R*_0_ is the maximum allowed extension of the bond. The parameters, *σ*, and *ϵ*_*f*_ represent the diameter of the bead and the strength of the interaction, respectively. The non-bonded polymer beads interact through the Lennard-Jones (LJ) potential given as

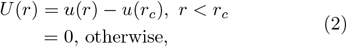

where 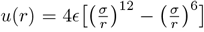. The cut-off distance *r*_*c*_ and the interaction strength *ϵ* are tuned to implement the philic and phobic nature of the polymer segments. We choose *r*_*c*_ = 2.5*σ* and *ϵ* = 2*k*_*B*_*T* for the attractive interaction between the non-bonded phobic beads, and *r*_*c*_ = 2_1*/*6_*σ* and *ϵ* = *k*_*B*_*T* for the pure repulsive interaction between non-bonded philic beads or between philic and phobic beads.

Thermal relaxation is introduced into the simulation through the following Monte Carlo moves. A bead is picked at random, and an attempt is made to move it to a new position within a box of size *l*. The move is accepted using the Metropolis criteria [71]. One Monte Carlo Step (MCS) consists of *N*_*p*_ attempts to move a bead. Henceforth, all time scales in this paper are expressed in the units of MCS. To change the state of a segment, one segment is picked at random, and the value of *ϕ* is changed from ±1 to ∓1 at a certain rate. Within one MCS, *R*_*f*_ ×*N*_*p*_ attempts are made to switch the state of the segments, which sets the timescale for the switching. Note that the segment locations along the polymer are clearly defined. For a polymer with *N*_*s*_ = 4, bead locations (1, 2, 3, 4) represent the first segment, locations (5, 6, 7, 8) represent the second segment, and so on. When the switching happens, the whole segment collectively switches as a single unit. Figure 1(a) shows a schematic picture of a copolymer consisting of both *ϕ* = 1 and *ϕ* = −1 beads for *N*_*s*_ = 4. Each polymer segment independently switches their state from *ϕ* = 1 to *ϕ* = −1 and vice versa with respective transition rates *R*_±_ and *R*_∓_, in units of 1*/*MCS, defined as,

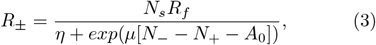

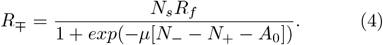

The corresponding bead (or segment) transition probabilities from *ϕ* = 1 to *ϕ* = −1 is *P*_+−_ = *min*(1, *R*_+−_) and *ϕ* = −1 to *ϕ* = 1 is *P*_−+_ = *min*(1, *R*_−+_). This transition probability depends on the switching attempt rate *R*_*f*_, instantaneous number of philic *N*_+_ and phobic *N*_−_ beads, asymmetry parameter 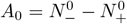, and a parameter *μ* that controls the fluctuations in *N*_+_ and *N*_−_. Here, 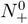 and 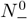 are the steady-state average value of *N*_+_ and *N*_−_, respectively. For the above two equations, we set 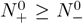, and the parameter 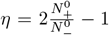, en-suring that *N*_+_ (*N*_−_) approaches the required values 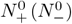 at steady state. When a polymer contains philic and phobic beads with 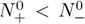, we need to make a change in the equations 3, 4 by replacing *η* in the denominator of eq 3 with 1 and the 1 in the denominator of eq 4 with 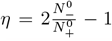. The cycle consisting of switching of segmental states and its thermal relaxation is inherently nonequilibrium, as demonstrated through the Kolmogorov loop in the supplementary material (see Fig. S1).

Based on equations 3 and 4, at steady state, the probability that the segments switch from philic/phobic to phobic/philic is 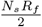. For a given switching attempt rate *R*_*f*_ and segment length *N*_*s*_, the corresponding polymer switching time can be written as 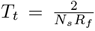, which is the time in which every polymer segment would have switched once on the average. Switching from one state to another is not allowed for the passive copolymer, i.e., *R*_*f*_ = 0 or *T*_*t*_ = ∞.

### Simulation Parameters

All the lengths are expressed in units of the diameter of the bead, *σ*. Energy is measured in the units of *k*_*B*_*T*. Time is measured in the units of MCS. The size of the polymer is *N*_*p*_ = 1024. The FENE parameters are 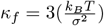, *ϵ_f_* = 0.305*k_B_T* and *R*_0_ = 1.3*σ*. The Monte Carlo bead moves are chosen within a box of size 0.2 × 0.2 × 0.2 centered at the bead. To explore the effect of switching rates, we consider a wide range of *R*_*f*_ from 10^−8^ to 10^−1^. All the results presented in this paper are averaged over at least 2400 independent configurations. Unless otherwise mentioned, configurations are sampled from MCS 4*τ*_*p*_ to 10*τ*_*p*_.

## III. RESULTS

We simulate chromatin as a diblock copolymer of length *N*_*p*_ having segments (blocks) of length *N*_*s*_, and study its 3D organization accounting for the nonequilibrium switching of chemical states of various segments in the polymer. As the polymer samples the configuration space, the chemical states of the segments are switched, between solvo-philic and solvo-phobic states, with the switching time *T*_*t*_. See section II for the model details.

We quantify the 3D organization by computing quantities like the radius of gyration, contact probability, and cluster size distribution at steady-state. The organization of symmetric copolymers (on average, the same number of philic and phobic segments, randomly distributed) of various segment lengths (*N*_*s*_ = 4, 8, or 16) are simulated using nonequilibrium Monte Carlo methods (see section II).

### A. Polymer compaction displays a non-monotonic behavior as a function of switching time

One of the ways to quantify the compaction of a polymer is to compute the radius of gyration, defined as [69]

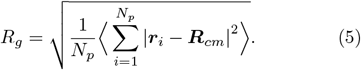

Here, ***r***_*i*_ is the position of *i*^*th*^ bead, and ***R***_*cm*_ is the center of mass position of the polymer. The angular brackets in equation 5 refer to the average over an ensemble of steady-state polymer configurations. *R*_*g*_ as a function of switching time *T*_*t*_ for different segment lengths (*N*_*s*_) are plotted in Fig. 2(a). We observe a nontrivial non-monotonic variation in *R*_*g*_ for all segment lengths, *N*_*s*_ = 4, 8, and 16. *R*_*g*_ has a well defined minimum between *T*_*t*_ = 10^5^ and *T*_*t*_ = 10^7^ and a maximum between *T*_*t*_ = 10^2^ and *T*_*t*_ = 10^3^. The switching time corresponding to maximum *R*_*g*_ increases with increasing *N*_*s*_. Moreover, for any *T*_*t*_, the radius of gyration increases with *N*_*s*_, though the polymer length and the fraction of philic to phobic beads remain the same. Below, we examine some steady-state configurations to understand this behavior of *R*_*g*_ as a function of *T*_*t*_.

**FIG. 2.**
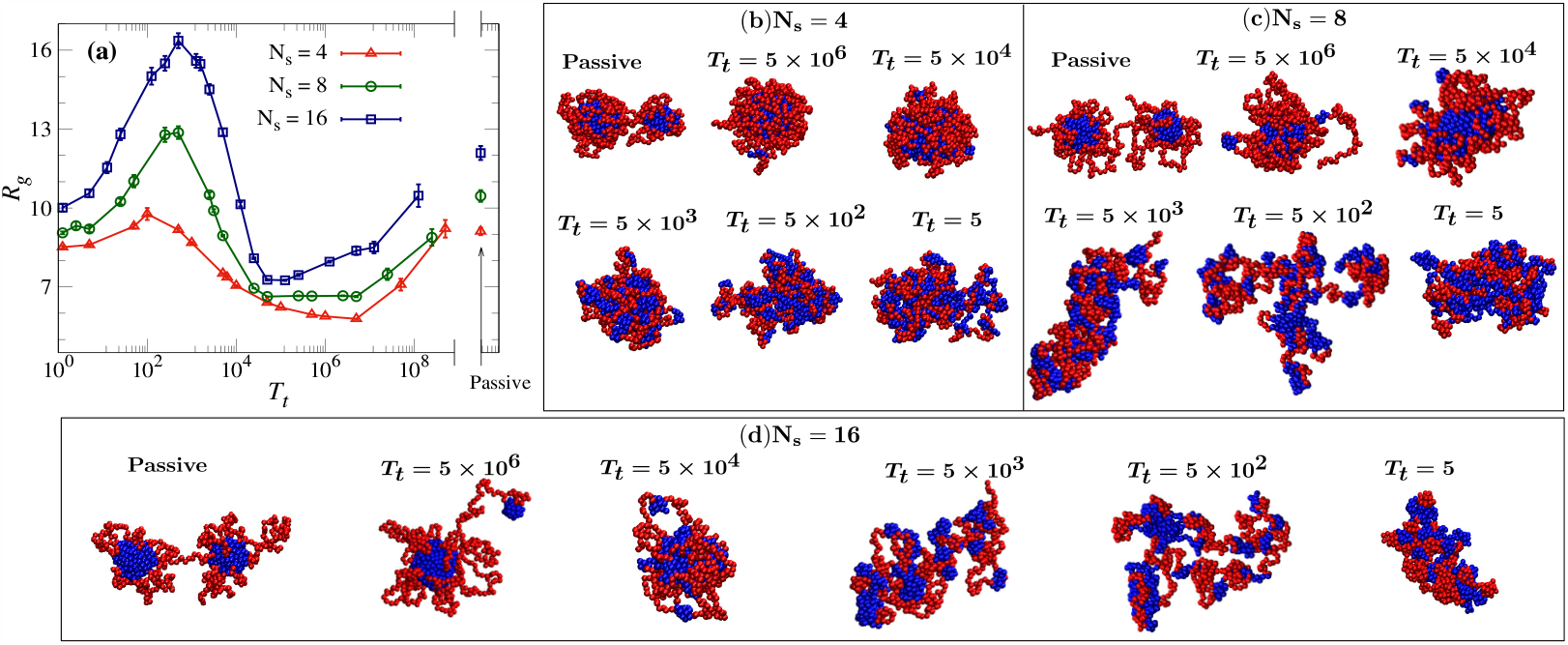
(a) Radius of gyration *R*_*g*_ of the polymer as a function of switching time *T*_*t*_ for different segment lengths *N*_*s*_. The *R*_*g*_ has maximum (minimum) compaction when *T*_*t*_ is of the order of the segment (polymer) relaxation time. *R*_*g*_ values of a passive polymer are included for comparison. (b)-(d) Steady state snapshots of the polymer for different switching time *T*_*t*_ and segment length *N*_*s*_. See the supplementary movie which captures the role of transition time on the polymer compaction and internal organization.

As shown in Fig. 2(b), for *N*_*s*_ = 4, we observe compact structures for intermediate transition times 5 × 10^6^ ≥ *T*_*t*_ ≥ 5 × 10^3^. This corresponds to the minimum in *R*_*g*_ (see Fig. 2(a)). For *T*_*t*_ ≥ 5 × 10^4^, including the passive situation, the structure appears to be segregated into a core-shell structure, with the phobic (blue) beads at the core surrounded by a shell composed of philic beads (red). Such morphology is absent when the segments switch their states faster (see configurations for *T*_*t*_ ≤ 5 × 10^3^ in Fig. 2(b)). Similar patterns are also observed for *N*_*s*_ = 8 and 16, although with more discernible differences in polymer compaction and shape (Fig. 2(c) and Fig. 2(d)). The shell is particularly prominent for *N*_*s*_ = 16 with the hair-like extensions of philic beads. Such hairy structures of philic segments at the surface are primarily responsible for the higher *R*_*g*_ with increasing *N*_*s*_ (see Fig. 2). These snapshots, which support the *R*_*g*_ observations, demonstrate that the steady-state polymer compaction depends on both the transition time and the segment length.

Next, to understand the locations of the minimum and the maximum values of *R*_*g*_ for different *N*_*s*_, we examine different natural timescales associated with the polymer. The relaxation time of the passive polymer *τ*_*p*_ is calculated from the correlation, 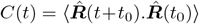, where, 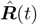 is the unit end to end vector. For *N*_*p*_ = 1024, *τ*_*p*_ is found be approximately 2 × 10^7^ MCS. Similarly, the relaxation time of a segment of size *N*_*s*_ is calculated by considering it as a short polymer. Since the polymer consists of both philic and phobic segments, whose relaxation time can be different, we define the segment relaxation time as *τ*_*s*_ = 0.5 × (*τ*_*ϕ*=1_ +*τ*_*ϕ*=−1_), wherein *τ*_*ϕ*=1_ and *τ*_*ϕ*=−1_ are the relaxation times of a philic and phobic segments, respectively. For *N*_*s*_ = 4, the segment relaxation time *τ*_*s*_ is approximately 300*MCS* and it increases as 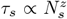, where *z* ≈ 2.

Our results suggest that *R*_*g*_ is at a minimum when the transition time *T*_*t*_ is of the order of the relaxation time *τ*_*p*_ of the polymer. This implies that polymer segments have enough time to interact and find a favorable configuration before any segment state is switched. Note that this minimum value of *R*_*g*_ is below the *R*_*g*_ obtained for the passive polymer. This indicates the presence of deep local minima in the configurational space of the passive polymer resulting in multiple clusters (see Fig. 2(b)-(d) for configurations). Switching of segmental states anneals the system by taking it out of these local minima and thus allowing it to explore all configurations. We will come back to this point later in the manuscript while discussing the contact probabilities.

When the switching time *T*_*t*_ is comparable to the segment relaxation time *τ*_*s*_, we see a maximum in *R*_*g*_ (Fig. 2(a)). Here, since the switching is quicker than the polymer relaxation time, distal phobic segments do not get time to come together and condense before the segments are switched. However, beads within each segment still get sufficient time to interact with each other. As they switch between phobic and philic states, each segment is physically collapsing or swelling. However, the timescales for swelling and collapse of the segments have an asymmetry, with the former being significantly higher than the latter (see supplementary material, Fig. S2). Since every segment goes through a phobic-philic transition within a time of *T*_*t*_, they all are in a partially collapsed configuration, leading to a state where the whole polymer looks like a self-avoiding string of (*N*_*p*_*/N*_*s*_) collapsed blobs (see Fig. 2(b)-(d)) and supplementary movie.

When the switching is faster (i.e., *T*_*t*_ *<< τ*_*s*_), the segments partially lose their identity as phobic or philic, and every pair of beads will now have a uniform effective attractive interaction leading to a decrease in *R*_*g*_. The string of blobs structure is now lost. However, the *R*_*g*_ still shows a dependence on the segment size arising from the asymmetry in transition times to and from the phobic state.

### B. Contact probability

Contact probability is one of the highly useful and experimentally measurable quantities to understand chromatin polymer organization. It measures how often different parts of the genome come in “contact”, given an ensemble of chromatin structures generated from a population of cells. This quantity helps us to understand the physical nature of chromatin organization and provides vital clues to its function [11, 25]. Contact probability is defined as follows: if the 3D spatial distance (*r*) between two beads is ≤1.3, then we consider those beads to be in contact at that instant. Given two beads *i* and *j* separated by a distance *S* = |*i* − *j*| along the polymer contour (known as genomic distance or chemical distance), the contact probability is defined as [72]:

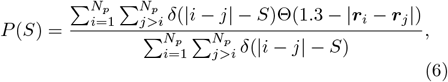

where Θ is Heaviside step function with Θ(*x*) = 1 for *x* ≥ 0 and zero otherwise. In general, the contact probability follows a relation *P*(*S*) ∝ *S*^−*γ*^ where the exponent *γ* depends on the nature of the polymer. For the self-avoiding walk (SAW), the contact probability exhibits a relation *γ* ≈2.18 [73]. In the case of polymers in a poor solvent, the equilibrium globular state exhibits *γ* = 1.5 for 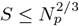 and *P*(*S*) is *constant* for 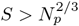 [74, 75], and for a fractal globule, *P*(*S*) ≈ *S*^−1^ [25, 74].

*P*(*S*) obtained from our simulations, with chromatin modeled as a diblock copolymer that actively switches the segment between the *ϕ* = 1 and *ϕ* = −1 states, for various switching times and segment lengths are presented in Figures 3(a)-(c). For small values of *S*, we see two distinct regimes, fast decay corresponding to the organization within a segment *S < N*_*s*_ and slow decay for 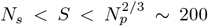. For the passive polymer (*T*_*t*_ = ∞), the slow decrease in *P*(*S*) is followed by a rapid decay beyond *S* ∼ 200. This behavior of the contact probability indicates the local collapse of the polymer to form globules of the size of approximately 200 and the formation of a superstructure resulting from the aggregation of these globules/domains/clusters, as shown in Fig. 4(a). This is similar to what is reported in [53].

**FIG. 3.**
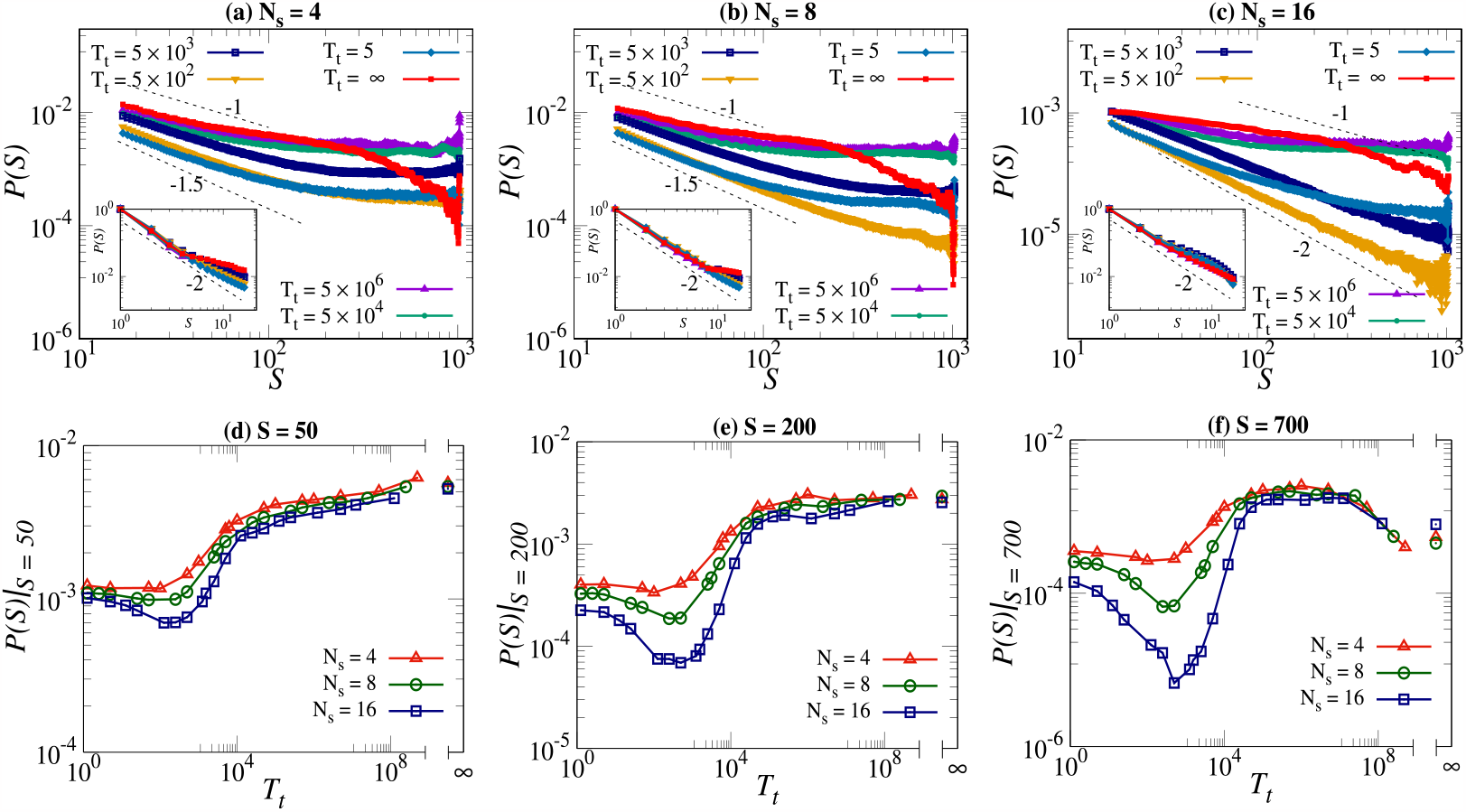
(a)-(c) Contact probability, *P*(*S*) as a function of genomic/chemical distance along the polymer (*S*). The inset shows the behavior of *P*(*S*) with *S* for *S* ≤ 16. The dashed lines represent different theoretical exponents to guide the eye. (d)-(f) Contact probability as a function of transition time for a fixed separation along the contour for various segment sizes. The respective genomic separation (S) is mentioned in every figure.

**FIG. 4.**
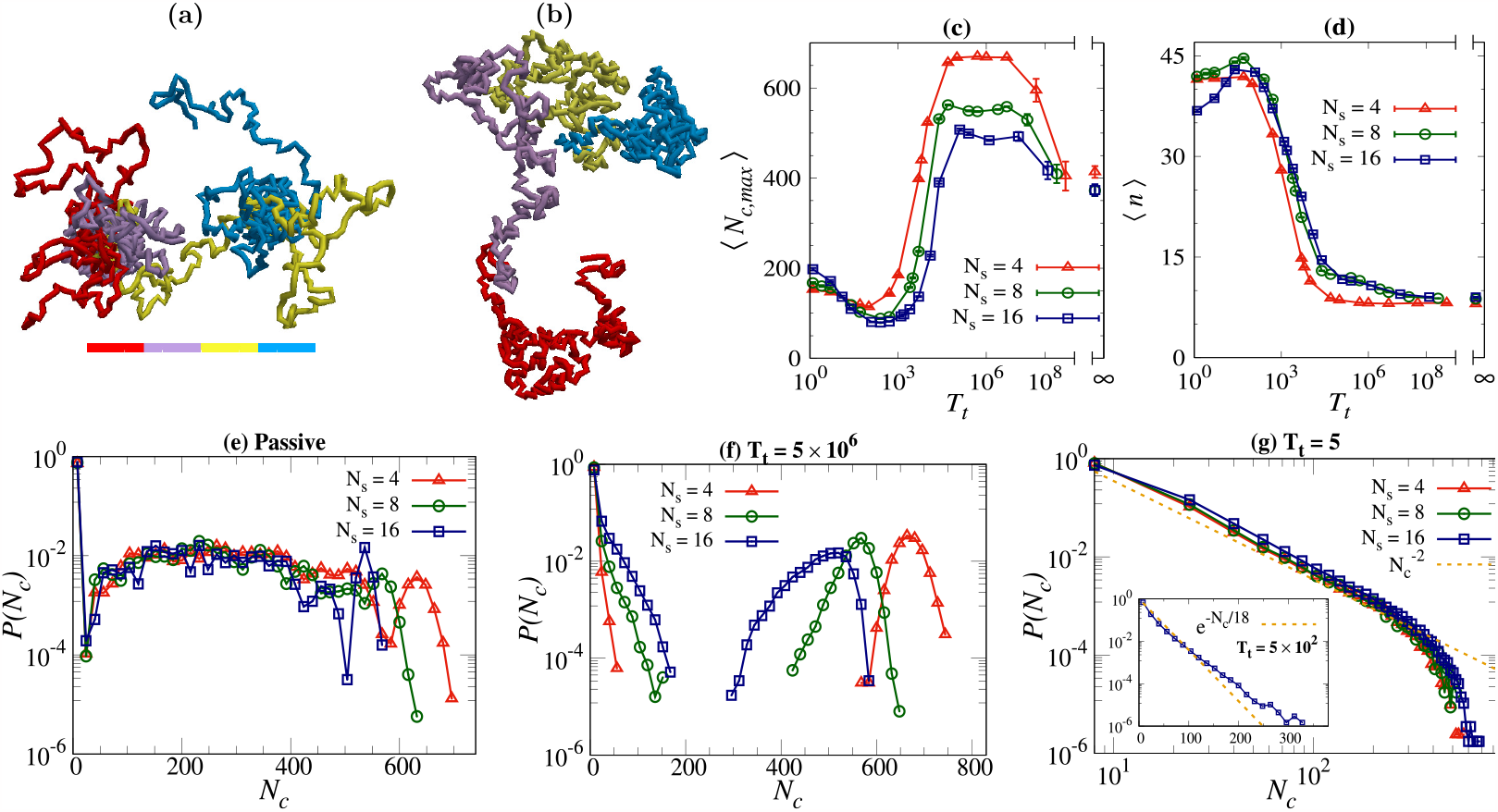
(a, b) Snapshots showing clustering of the polymer for *N*_*s*_ = 16 at *T*_*t*_ = ∞ (passive) and *T*_*t*_ = 10 respectively. The color-bar indicates the sequence of colors used to mark different parts of both polymers. (c) The average size of the biggest cluster ⟨*N*_*c,max*_⟩ as a function of *T*_*t*_ for different *N*_*s*_ values. (d) Variation of average cluster number, ⟨*n*⟩ with respect to *T*_*t*_ for different segment lengths. Probability distribution of cluster size *P* (*N*_*c*_) for (e) passive polymer, (f) *T*_*t*_ = 5 × 10^6^, and (g) *T*_*t*_ = 5 with inset for *T*_*t*_ = 5 × 10^2^. The dashed lines in (g) show the power/exponential fit.

For the polymers with finite segment switching time, for *S < N*_*s*_, the contact probability remains the same as in the passive case. However, for *S > N*_*s*_, the generic behavior, in the presence of switching, is a power law decrease in *P*(*S*) approaching a constant high value. The power law exponent and the value at which *P*(*S*) saturate depend on *N*_*s*_. The saturation of *P*(*S*) is most prominent, and above the value for passive polymers, when the switching is in the timescale of polymer relaxation. This suggests that segment state switching at these rates helps in increasing the mobility of the segments and organizes the polymer into a steady-state configuration wherein the average distance between any two points is independent of their genomic separation (see supplementary Fig. S3 for 3D distance plots). The value of *S* at which *P*(*S*) saturates depends on *N*_*s*_ and *T*_*t*_. For *N*_*s*_ = 4, as shown in Fig. 3(a), *P*(*S*) saturates for all values of *T*_*t*_, with the saturation point shifting to higher values of *S* with increasing *T*_*t*_. As *N*_*s*_ increases, Fig. 3(a)-(c), for any fixed values of *T*_*t*_, the saturation point is shifted to a higher value of *S*. It’s clear from the above that the steady-state structures of such transition active polymers are neither fractal nor equilibrium globule.

Figure 3(d)-(f) depicts the contact probability as a function of *T*_*t*_ for a fixed genomic separation for different segment lengths. It is seen that the contact probability for a particular genomic separation changes non-monotonically with switching time. This is increasingly apparent for larger *N*_*s*_. Another intriguing point to note is that the contact probability decreases with increasing segment length. Also, for *N*_*s*_ *>* 4, *P*(*S*) exhibits minima when the switching time is comparable to the segment relaxation time *τ*_*s*_. The effect of switching time on the contact probability is significant for *S >> N*_*s*_ with a broad maxima for large *S*, around *T*_*t*_ ∼*τ*_*p*_, corresponding to the minima in *R*_*g*_, see Fig. 3(f) and Fig. 2(a). Thus for large genomic distance *S >> N*_*s*_, the contact probability is independent of *S* and is maximum when the switching time is comparable to *τ*_*p*_. The effect of switching time on the contact probability is more significant for *S >> N*_*s*_.

### C. Cluster size distribution

In the previous sections, we discussed the non-monotonic behavior of the radius of gyration and the contact probability of the polymer with respect to the transition time. The non-monotonic nature of the radius of gyration implies that for multiple values of the transition times (*T*_*t*_), the numerical value of *R*_*g*_ could be the same. However, the snapshots in Fig. 2(b)-(d) suggest the conformation of the polymer is very different even when their *R*_*g*_ values are the same, essentially implying different arrangements of philic and phobic beads along the polymer. Thus quantifying compaction through *R*_*g*_ alone is not sufficient to describe the system.

To further investigate the structure of the polymer at different values of *T*_*t*_, here we analyze the clusters present in the polymeric system for different *T*_*t*_ and *N*_*s*_ in the following way. If any two non-connected beads are separated by a 3D distance *<* 1.3, they are considered part of the same cluster. A cluster can have both philic and phobic beads in it. Cluster size *N*_*c*_ is defined as the number of beads in a given cluster. In the analysis, we only consider clusters having *N*_*c*_ ≥ 4, the size of the shortest segment considered here.

The average size of the biggest cluster ⟨*N*_*c,max*_⟩ and the average number of clusters ⟨*n*⟩ at steady state are plotted in Fig. 4(c),(d). ⟨*N*_*c,max*_⟩ reflects the non-monotonic behavior of *R*_*g*_, with a minimum when *T*_*t*_ ≈ *τ*_*s*_ and a broad peak when *T*_*t*_ ≈ *τ*_*p*_. Based on the understanding we gathered in the above sections through analyzing the configurations, *R*_*g*_, and *P*(*S*), we expect a SAW of clusters of size *N*_*s*_ (as shown in Fig. 4(b)) when *T*_*t*_ ≈ *τ*_*s*_. Consequently, the number of clusters is the largest, and the size of the biggest cluster is the least at these switching times. When switching is faster (*T*_*t*_ *< τ*_*s*_), the beads interact with an effective marginal attraction, resulting in a small increase in the cluster size. On the other hand, as *T*_*t*_ approaches the polymer relaxation time (*τ*_*p*_), more and more regions of the polymer interact, the number of clusters decreases, and the cluster size becomes the largest. A comparison between Fig. 2(a) and Fig. 4(c) tells us that the polymers, though may have similar *R*_*g*_ values, have very different internal structures at different *T*_*t*_ values.

To understand the clustering of the segments further, we compute *P* (*N*_*c*_) — the probability of having a cluster of size *N*_*c*_ — for different *T*_*t*_. Figure 4(e)-(g) captures the distinguishable variations in *P* (*N*_*c*_). The non-zero values of *P* (*N*_*c*_) for 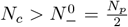 indicate that the cluster is a mixture of philic and phobic beads. As illustrated in Fig. 4(e), for the passive polymer (*T*_*t*_ = ∞), there are two noticeable peaks, a broad one from *N*_*c*_ = 100 to 400 and a sharper one at a value of 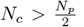. The broad peak arises due to the long-lived metastable states, which for the size of the polymer considered here, mainly consists of two distinct clusters of varying sizes. The peak at a higher value of *N*_*c*_ represents configuration with a single big cluster. The position of this peak shifts to larger values of *N*_*c*_ with decreasing segment size *N*_*s*_. Such a shift of peak to the right is due to an increase in the mixing of philic and phobic beads with decreasing *N*_*s*_, as can be seen from Fig. 2(b)-(d).

Figure 4(f) depicts the cluster distribution when the switching time is comparable to the polymer relaxation time. The distribution has one prominent peak at 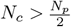. This is consistent with our understanding of large average cluster size and high compaction (small *R*_*g*_). The shifting of the peak towards the left with increasing *N*_*s*_ is due to less mixing of philic and phobic beads with segment length. It is interesting to note the absence of medium-sized clusters; this suggests that the interplay between switching and polymer relaxation has led to the coalescence of medium-sized clusters into larger clusters.

When the switching time is very short (*T*_*t*_ *< τ*_*s*_), we obtain power-law distributions in cluster sizes (see Fig. 4(g)), indicating the absence of any length scales for cluster sizes. For *T*_*t*_ ≈ *τ*_*s*_, *P* (*N*_*c*_) decays exponentially, and the associated length scale is approximately equal to the segment length (see inset of Fig. 4(g)), consistent with the segment level clustering discussed earlier.

### D. Polymer organization and mixing of philic and phobic beads

We look into the local packing fractions inside a cluster to examine the positional arrangement of philic and phobic beads. The local packing fraction of philic/phobic beads within a cluster is defined as 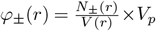, in a spherical shell of volume *V* (*r*) between *r* to *r*+*δr*. Here, *r* is the distance of a bead from the center of mass position of its cluster, and *V*_*p*_ is the volume of a single bead 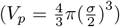. For both *φ*_+_(*r*) and *φ*_−_(*r*) calculations, we average over only big clusters with size 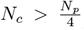. Figures 5 (a)-(d) show variation in *φ*_+_(*r*), *φ*_−_(*r*), and *φ*_+_(*r*) + *φ*_−_(*r*), as a function of *r/R*_*gc*_ as a function of *T*_*t*_. Here, *R*_*gc*_ is the radius of gyration of the considered cluster. We choose *N*_*s*_ = 16 for these calculations. For the passive polymer, when *r/R*_*gc*_ ≤ 1, *φ*_−_(*r*) is very large and *φ*_+_(*r*) is nearly absent, implying a cluster core that is only made up of phobic beads. For large *r/R*_*gc*_, *φ*_+_(*r*) dominates over *φ*_−_(*r*) indicating the presence of philic beads near the surface of the cluster. This behavior of *φ*(*r*) is consistent with the snapshots shown in Fig. 2.

**FIG. 5.**
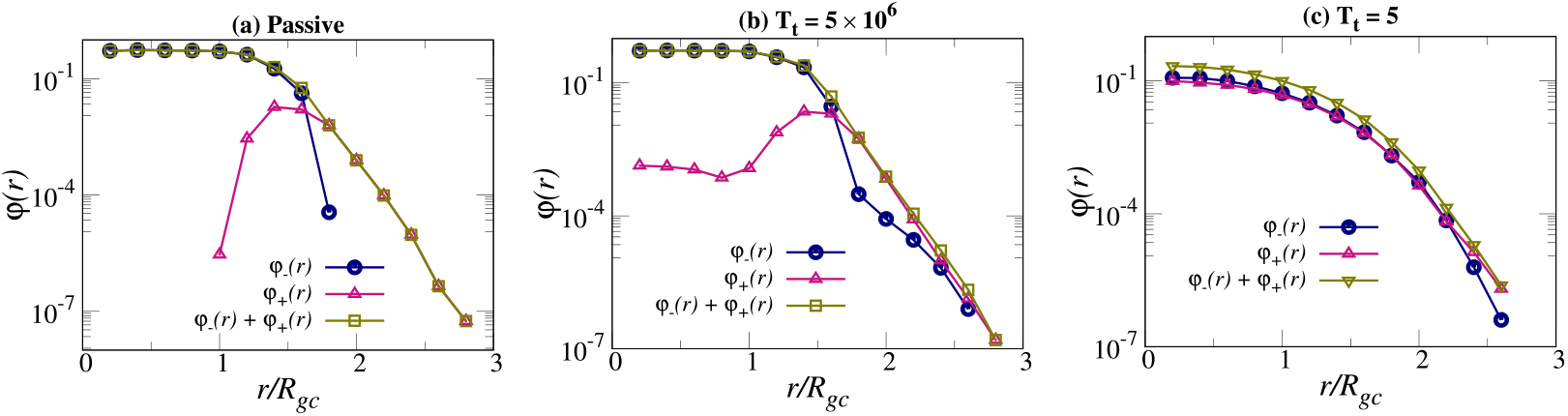
(a)-(d) Local packing fraction of philic *φ*_+_(*r*), phobic *φ*_−_(*r*), and all beads in a cluster for different *T*_*t*_ with *N*_*s*_ = 16.

When switching is introduced, the philic and phobic beads slowly start mixing. For *T*_*t*_ = 5 × 10^6^, *φ*_−_(*r*) *> φ*_+_(*r*), for *r/R*_*gc*_ ≤ 1 (Fig. 5 (b)). This implies that the interior of the cluster is still dominated by the phobic beads with the presence of a small number of philic beads. For *r/R*_*gc*_ *>* 1, much more mixing has happened and we find *φ*_−_(*r*)≈ *φ*_+_(*r*). As shown in Fig. 5(c), for very fast switching, there is a complete mixing of philic and phobic beads, and the difference between volume fractions vanishes.

## IV. DISCUSSION

Regions of chromatin are known to switch between protein-bound/unbound or active/repressed states. These transitions happen at different length scales, typically from a nucleosome to a gene. In this paper, we presented a copolymer model that allows for chemical reactions to switch the segmental states between solvophilic (active) and solvo-phobic (repressed) states in a nonequilibrium way. This nonequilibrium switching is achieved by ensuring that transitions follow a fixed rate and thus do not obey detailed balance [59] (see supplementary material for more details). By computing the radius of gyration, cluster size distribution, and contact probabilities as a function of the switching rate, we examined how such nonequilibrium switching could affect the chromatin organization. Though diblock copolymer models with philic/phobic segments were considered earlier, the active switching between these two states was not explored. We show that such switching has a nontrivial effect on the 3D organization of the polymer and may play a role in determining the structure of the chromatin in addition to other protein-mediated looping and condensation.

Two timescales inherent to the block copolymers are the relaxation time of the segment (*τ*_*s*_) and the relaxation time of the polymer (*τ*_*p*_). As can be seen from Fig. 2, a sweep of the transition time *T*_*t*_ through these timescales results in a significant change in the steady-state configuration of the polymer, exhibiting a non-monotonic change in compaction. We will discuss two important regimes wherein *T*_*t*_ ∼ *τ*_*p*_ and *T*_*t*_ ∼ *τ*_*s*_.

A diblock copolymer is known to have multiple deep minima in its configuration space [76, 77], and a passive polymer can get stuck in one of these long-lived metastable states. Our results show that switching the segment states at a low rate is enough to allow the polymer to come out of such metastable states and better explore the configuration space, also shown in Fig. 2. This is reflected in the decrease of *R*_*g*_, compared to its value for the passive polymer, as the switching is turned on at a low rate, reaching a minimum when *T*_*t*_ ≈ *τ*_*p*_. The higher compaction and dynamics of the clusters lead to an increase in contact probabilities between regions with higher genomic separation. This has relevance, especially considering that solvo-phobic/philic regions can be mapped as nucleosome bound/unbound regions, to recent experiments which suggest that far away regions of chromatin come in proximity even in the absence of loop extruders [30, 78]. Specifically, far-away genes are known to come together to form compartments and transcription factories [79]. In a passive polymer, the probability of coming together of regions spaced far away along the contour decreases as a function of contour separation. This arises due to the hierarchical clustering of the copolymer [53]. Our results suggest that slow switching (*T*_*t*_ ∼ *τ*_*p*_) allows the clusters to reconfigure and thus makes the contact probability of far-away segments independent of the contour separation. This is also highlighted in the spatial distance matrix shown in the supplementary material, Fig. S4. This physical phenomenon is likely to help the formation of compartments as well as transcription factories. At the small switching rates (high *T*_*t*_), there is only one dominating cluster, which is a mixture of phobic and philic beads (see Fig. (4,5)). The amount of mixing depends on the segment length. More mixing and larger cluster sizes are observed for smaller segment lengths.

The polymer exhibits a completely different behavior when the switching time *T*_*t*_ is small and comparable to the average segment relaxation time *τ*_*s*_. As seen in Fig. 2(a), the polymer is now in its maximum swollen state. The corresponding configurations, shown in Fig. 2(b)-(d), indicate a string of clusters. This is due to the asymmetry in the segment relaxation time going from the collapsed state to the swollen state and vice versa. The former time being larger, the segments tend to retain the collapsed configuration when *T*_*t*_ is smaller than the relaxation time for the philic segments (see supplementary Fig. S2). As a result, we see a self-avoiding walk of blobs at the segmental level, as shown in Fig. 4(b). The corresponding cluster distribution is an exponential function, indicating the existence of a segment-dependent typical cluster size (see Fig. 4). The contact probability at this switching rate is the lowest for genomic separation larger than the segment length. When the switching rate is increased further, beyond the segmental relaxation time, the segments switch before they can thermally relax. As a result, the polymer behaves like a self-avoiding walk of a homogeneous string with an average uniform attractive interaction between beads. The resulting cluster distribution is a power law. It is intriguing that, in this regime, *R*_*g*_ decreases with the switching rate.

If one considers a small segment as a nucleosome and the cluster as a TAD, our results shown in Fig. 5 suggest that the switching (nucleosome on/off dynamics) could be thought of as a regulatory mechanism to control accessibility; this can bring any segment to the exterior of the TAD and make them accessible to diffusing enzymes/proteins.

There have been many discussions about the gel versus fluid nature of the chromatin domains [80–82]. In this context, to understand the dynamic nature of the clusters formed at various switching rates, we have computed the self part of the van Hove function *G*_*s*_ [83] in the polymer center of mass frame and mean square displacement (MSD) averaged over of all beads. While the peak in *G*_*s*_ is a measure of the average mobility of the beads, its width is a reflection of compaction of the polymer. The MSD shown in Fig. 6(a) demonstrates that the mobility of the beads is sub-diffusive and, like the *R*_*g*_ of the polymer, is a non-monotonic function of the switching time. It is clear from Fig. 6(b) that the clusters are fluid-like with bead mobility that depends on *T*_*t*_.

**FIG. 6.**
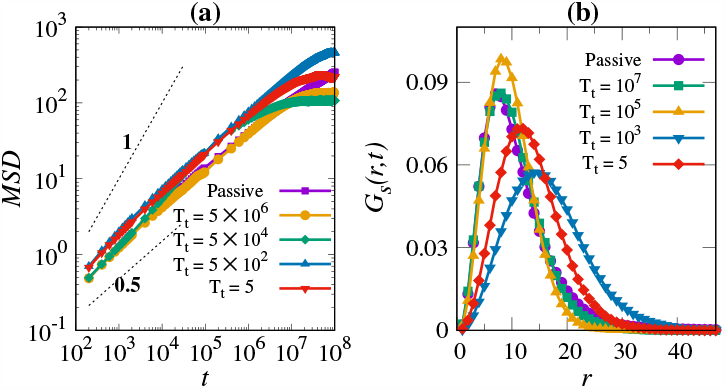
(a) Self-part of the van Hove function as a function of *r* for different values of switching time, at *t* = 10^7^. The dashed lines represent different exponents to guide the eye. (b) The mean square displacement of the polymer beads in the polymer center of mass frame, averaged over all beads.

In summary, we have focused on the effect of nonequilibrium switching of the segments in a copolymer model for chromatin and studied its influence on the organization of the whole polymer. We have demonstrated that such inherently nonequilibrium activity can alter the contact probability and compaction of the polymer. It is important to note that the nuclear environment is highly heterogeneous, and the relaxation time of the segments can vary across regions from a few seconds to a few minutes [84]. Thus for a given rate of segmental modification, the response of the polymer could vary across regions. The generic model presented here can be modified to include these additional complexities to obtain a location-specific contact map.

## ACKNOWLEDGMENTS

We acknowledge the National Supercomputing Mission grant DST/NSM/R&D HPC Applications/2021/03.43. The support and the resources provided by ‘PARAM Brahma Facility’ under the National Supercomputing Mission, Government of India, at the Indian Institute of Science Education and Research (IISER) Pune are gratefully acknowledged. We would like to acknowledge the HPC facility at IIT Palakkad (Chandra cluster) for performing some computational work. S. Kadam acknowledges CSIR, Govt. of India, for Senior Research Fellowship.

