## Supplementary information for "Nonequilibrium switching of segmental states can influence compaction of chromatin"

#### 1 Nonequilibrium nature of switching and Kolmogorov loop

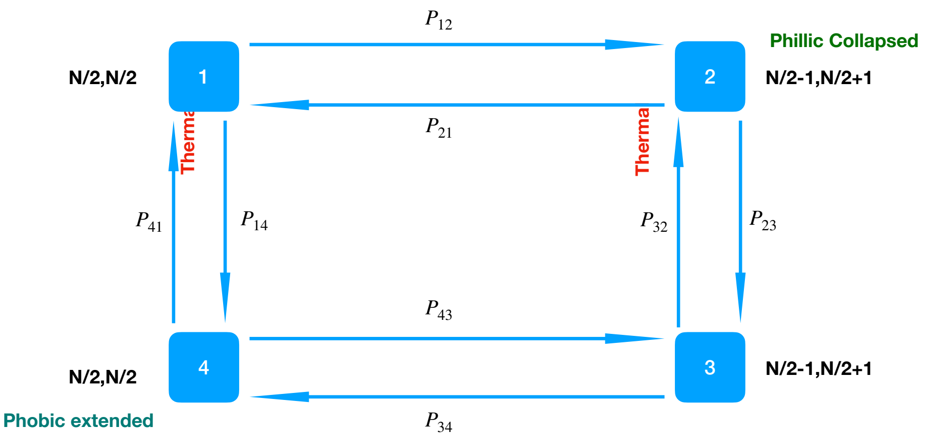


**FIG. S1:** The Kolmogorov loop demonstrating nonequilibrium nature of switching between philic and phobic states (see SI text).

To illustrate the nonequilibrium nature of the switching transitions, we show, in Fig. S1, the Kolmogorov loop for the transition of the segments when the number of phobic and philic segments are equal on average. State 1 is where both the phobic and philic segments are in equilibrium. In state 2, one of the phobic segments is switched to a philic state. This "excited state" configuration relaxes to its equilibrium segmental state at state 3. From state 3, one philic segment is switched to phobic, and this constitutes state 4.

Generalizing equations(3, 4), for different attempt rates between states, and with $N_+^0 = N_-^0 = N/2$, we can write the transition probabilities as $P_{12} = \frac{R_{12}}{2}, P_{23} = 1, P_{34} = \frac{R_{34}}{1+e^{-\mu}}, P_{41} = 1, P_{14} = e^{-\beta E_1}, P_{43} = \frac{R_{43}}{2}, P_{32} = e^{-\beta E_2}$, and $P_{21} = \frac{R_{21}}{1+e^{-\mu}}$, where $R_{ij}$ is the generalised attempt rate for transition from state-$i$ to state-$j$. Please note that the results presented in the manuscript are for $R_{ij} = R_f$ for all $(ij)$

The Kolmogorov condition for equilibrium is $P_{12}P_{23}P_{34}P_{41} = P_{14}P_{43}P_{32}P_{21}$. This implies, for equilibrium switching

$$R_{12}R_{34} = R_{43}R_{21}e^{-\beta(E_1+E_2)}. \tag{S1}$$

Here $E_1$ is the energy difference between the extended and collapsed configurations of a phobic segment. Similarly $E_2$ is the energy difference between the collapsed and extended configurations of a philic segment.

From the above arguments, it is clear that the switching of states is inherently nonequilibrium, and equilibrium switching can happen only in a very limited region in the $R_{ij}$ space. Equation S1 suggests that for switching to be in equilibrium, since $E_1 + E_2 > 0$, the attempt rates of switching from phobic segments in the extended state and philic segments in the collapsed state, should be much larger than the switching from the equilibrated segmental states, which is expected since these excited states are short-lived.

#### 2 Asymmetry in relaxation time of philic and phobic states

In the simulations reported here, the transition rates from phobic to philic and vice versa are the same. However, there is considerable asymmetry in the relaxation time of the segments depending on whether they are switched from phobic or philic state. We check this by calculating the auto-correlation $C(t)$ of the unit end to end vector of a homogeneous polymer of length 512, when switched from the equilibrated phobic/philic state to the philic/phobic state. This is shown in Fig. S2. It is clear from the figure that the relaxation is considerably faster when switch form philic to phobic state.

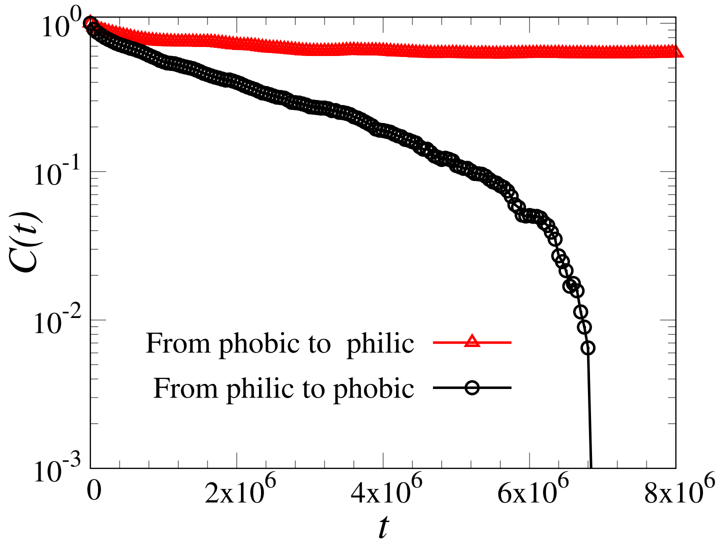


**FIG. S2:** Relaxation of unit end to end vector of a polymer of length 512 when switched from phobic to philic and vice versa.

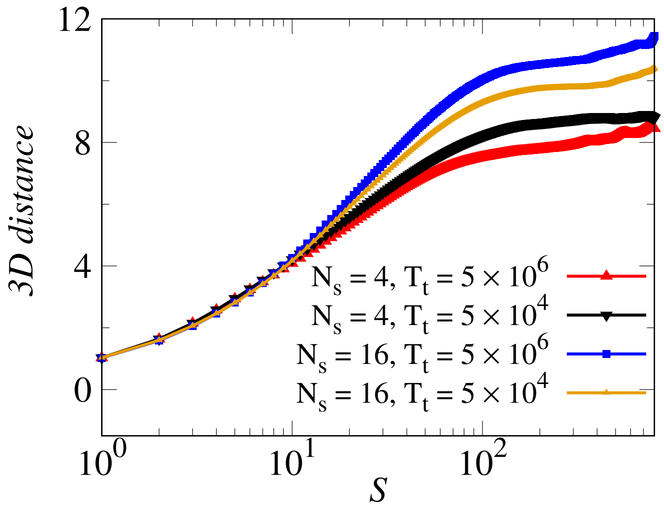


**FIG. S3:** Average 3D distance between pairs of beads of the polymer separated by genomic distance $S$ for different combinations of $N_s$ and $T_t$ values.

### 3 Spatial distance matrix

Figure S4 shows the spatial distance matrix for different switching times. For a passive polymer (see Fig. S4(a)), we see two TAD-like structures (domains) arising due to two compact clusters trapped in long-lived meta-stable states. Please note that these plots are generated from one realization after averaging over many steady-state time observations. Since there is no boundary-determining factors, the location of the boundary and size of the domains may vary in different realizations. In Fig. S4(b), where the polymer switching time is comparable to the polymer relaxation time, we see more interaction among the beads irrespective of their chemical nature and hence average distance between any two beads is small (a compact polymer). In Fig. S4(c), the corresponding structure is an SAW of blobs. This results in a locally folded polymer as reflected in the Fig. S4(c) (also see Fig. 4(b)).

### 4 Supplementary video - I

In this video we show the dynamics of a di-block copolymer of length 1024, with solvo-phillic and solvo-phobic segements of length 16, as the segments are switched from phobic/philic to philic/phobic and vice versa in a swithcing time of $T_t$. The switching time is varied from the fast switching ($T_t = 5$) to the case of the passive polymer ($T_t = \infty$). See main text for details of the model.

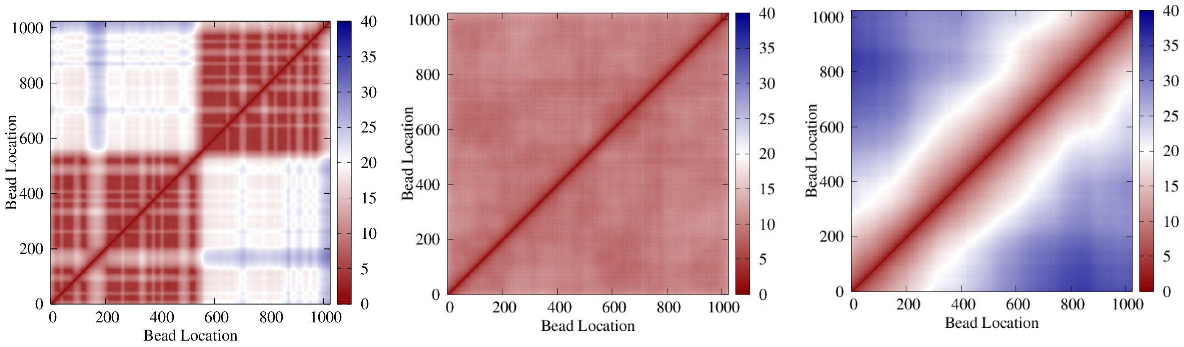


**FIG. S4:** Spatial distance matrix — 3D distance between all bead pairs between $i$ and $j$ — for a copolymer with solvo-phobic and solvo-philic segments of size $N_s = 16$. (a) Passive polymer with segments distributed randomly, (b) polymer with slow switching of the segmental states ($T_t \approx \tau_p$), and (c) polymer with fast switching ($T_t \approx \tau_s$). These plots are for a single realisation averaged over time.
